# Ethanol pre-exposure enhances alcohol-seeking behavior at cellular level by chemoattraction and exhibits bleb-driven cellular stress response in uniform ethanol concentration

**DOI:** 10.1101/2021.07.28.454082

**Authors:** Neelakshi Kar, Jayesh Bellare

## Abstract

This study investigates the effect of ethanol and its pre-exposure on cell migration. Here, fibroblast cells were first pre-treated with ethanol, and their migratory behavior was tested in both chemotaxis and chemokinesis setup. 1% ethanol was taken as a potential chemotactic agent. The study reveals that in presence of ethanol gradient cells display migration towards ethanol, and pre-exposure further augments this migratory behavior by altering their chemotactic responsiveness. In uniform ethanol concentration, cells first undergo three staged adaptations to the new environment: shrinking, blebbing, and recovery, where cells use bleb-driven cell protection machinery to adapt. Thus, migration is initially stalled. But once the cells resume locomotion, no significant difference in migratory parameters is observed. Overall, this study establishes ethanol as a chemoattractant for fibroblasts, with cells showing enhanced alcohol-seeking behavior upon pre-exposure. Such behavior is reminiscent of seeking and tolerance exhibited by alcohol-dependent addictive behavior in higher organisms including humans.

## 1. Introduction

Cell migration plays a very crucial role in the physiological well-being of multicellular organisms. Right from the development of the embryo to tissue repair and immune response, an orchestrated movement of cells is a prerequisite for the proper functioning of the biological system (Trepat et al., 2012). Anomalies pertaining to cell migration upon ethanol exposure have been studied in the last two decades. These reports indicate impairment of wound healing mechanism including reduced angiogenesis and re-epithelialization (Guo and DiPietro, 2010), developmental defects owing to the aberration of premature neuronal cell migration (Jiang et al., 2008; Kumada et al., 2007), and inhibition of L1 mediated cell-cell adhesion (Ramanathan et al., 1996). Ethanol has also been reported to promote metastasis of tumor cells by up-regulating the expression of Monocyte Chemoattractant Protein-1 (MCP-1) (Wang et al., 2012; Xu et al., 2016). Thus, ethanol exposure alters the migration ability of cells depending on cell type. However, ethanol sensing and migration to ethanol per se and cell motility in uniform ethanol concentration is not a well-studied area of research.

In the prokaryotic system, negative chemotaxis to ethanol has been reported in *Ralstonia pseudosolanacearum* Ps29 (Oku et al., 2017) and positive ethanol taxis in *B. subtilis* (Tohidifar et al., 2020). However, ethanol as a chemotactic agent has not been studied in eukaryotic cells to the best of the authors’ knowledge. Ethanol is an addictive drug, whose addictive property is attributed to complex neurobiological mechanisms, involving numerous neural circuits in different parts of the brain (Cui et al., 2013). But, can addictive drugs also act as attractants for cells? Moreover, can chemotaxis be used as a tool to trace down addiction to lower levels of the cellular organization? The cellular approach to addiction studies, though seemingly inconsequential, can help understand genetic differences that dictate addictive behavior (Crabbe, 2008). Moreover, the cell behavior in terms of motility in uniform ethanol concentration also presents an interesting facet. This is because cells first perceive ethanol as a stressor upon its introduction to the cellular environment, which leads to cell shrinkage, and migration is expected to be halted. Our previous study on cell behavior in uniform ethanol concentration indicated cell shrinkage and adaptability of fibroblasts in a dose-dependent manner (Kar et al., 2021). In this study, we look into the details of cell behavior and motility in uniform ethanol concentration.

This study uses chemotaxis as a tool to understand if ethanol-seeking behavior exists in the cellular system, and evaluates the cellular dynamics in the chemokinesis mode of migration. It investigates two different aspects of cell migration pertaining to ethanol: first, migration of cells in presence of ethanol (gradient as well as uniform concentration) and, second, the effect of ethanol pre-treatment on migration. The focus of this study is to evaluate the chemotactic potential of ethanol on the cellular system. In this study, NIH3T3 mouse fibroblast cells were first grown in presence of ethanol for seven days, and then their migration ability was analyzed in both chemotaxis and chemokinesis setup.

## 2. Materials and Methods

### 2.1. Materials

NIH 3T3 (Mouse Embryonic Fibroblast) cell line was purchased from NCCS, Pune, India. The cells were maintained in Dulbecco’s Modified Eagle’s Medium (DMEM) which was obtained from HiMedia, India. Other chemicals such as Fetal Bovine Serum (FBS), Trypsin-EDTA solution, Antibiotic Penicillin-Streptomycin, Crystal violet, and Trypan blue were also obtained from Himedia, India. Acetic acid was purchased from Merck, US. Glutamax was purchased from Thermo Fisher Scientific, US. Acetic acid was purchased from Merck, US. Fluorescein isothiocyanate-Phalloidin (FITC-Phalloidin) and 4’, 6-diamidino-2-phenylindole (DAPI), were obtained from ThermoFisher Scientific, US. FITC Annexin V Apoptosis Detection Kit I (BD Pharmingen™, Cat no. 556547) was obtained from BD Biosciences, US. Cell culture inserts of polyester (PET) membrane, 8 μm pore size, were purchased from BD Falcon. Ethanol used in this study is of HPLC grade from Commercial Alcohols, Greenfield Global, Canada. All the reagents and chemicals are of analytical grade.

### 2.2. Maintenance of cell lines and ethanol pre-exposure

NIH 3T3 cells were grown in T25 flasks in DMEM media with 10% FBS, and 1% antibiotic solution. The cells were maintained at 37ºC with 5% CO_2_ in an incubator (ThermoFisher Scientific, USA). Cells from 70-80% confluent flasks were used in experiments.

Two different dosing strategies were used for this study. Firstly, 2.5 × 10^3^ cells were seeded in six different wells of the 6-well plate. After 12 h of cell attachment, cells were exposed to ethanol in two different ways:

a. Acute-Periodic – Cells were exposed to 0.01%, 1% and 3% (v/v) ethanol concentrations for 1 h/day for 7 days. For further reference, these treatment groups have been denoted as 0.01A, 1A, and 3A respectively.
b. Chronic – Cells were continuously exposed to 0.01%, 1%, and 3% (v/v) ethanol concentrations for 7 days i.e. the cells were grown in media containing ethanol of different concentrations. These treatment groups have been denoted as 0.01C, 1C, and 3C respectively.

For better understanding, different treatment conditions and their notations are presented in Table 1. The term ‘acute-periodic’ and ‘chronic’ has been used to indicate the time of exposure.

**Table 1:** Nomenclature for pre-treatment groups.

| S.N. | EtOH Conc. % (v/v) | Dosing | Nomenclature |
| --- | --- | --- | --- |
| 1 | 0 | None | 0 |
| 2 | 0.01 | Acute-Periodic | 0.01A |
| 3 | 1 | Acute-Periodic | 1A |
| 4 | 3 | Acute-Periodic | 3A |
| 5 | 0.01 | Chronic | 0.01C |
| 6 | 1 | Chronic | 1C |
| 7 | 3 | Chronic | 3C |

Cells were then harvested by trypsinization and used for desired experiments. It should be noted that the spent cell culture media were replaced by fresh media with the desired ethanol concentration every day.

### 2.3. Cell Proliferation Assay

To check cell proliferation after seven days of treatment, cells adhered to the well plate were trypsinized and the number of cells was calculated using Improved Neubauer Haemocytometer (Rohem, India) by diluting with 0.4% trypan blue dye before counting.

### 2.4. Apoptosis Assay

This assay was done using FITC Annexin V Apoptosis Detection Kit I. For apoptosis assay, the supernatant obtained each day was stored in a separate 6-well plate and kept in an incubator, providing the same growth condition as the adhered cells. Later, after 7 days of treatment, cells were harvested from both plates. Cell suspensions including the supernatant were centrifuged at 2500 x g for 5 min. The cell pellets obtained from each treatment group were washed with PBS twice. Cell pellets were then suspended in 100 μl of Annexin V Binding Buffer. 0.2 μl of FITC Annexin V dye was added to the cell suspension and incubated at room temperature for 15 min in dark. 400 μl of Annexin V Binding Buffer was further added. PI was added 5 min before analysis. The cell suspension was analyzed via flow cytometry (BD FACSAria Flow Cytometer, BD Biosciences, US). The experiment was done according to the manufacturer’s instructions. Data analysis was done in FlowJo™ Software V10.0.8. (Becton, Dickinson and Company, U.S.A.).

### 2.5. Cell Adhesion Assay

5×10^4^ cells from each treatment group were seeded in a 96-well plate and were allowed to stand for 1 h at 37ºC, 5% CO_2_. After 1 h, the supernatant was removed and cells were fixed with 70% ethanol for 10 min. 100 μl of 0.2 % crystal violet was added to each well for cell staining for 15 min. Wells were washed with distilled water 3-4 times to remove the extra dye. 10% acetic acid was then added to each well for dye elution and absorbance was measured at 592 nm with a spectrophotometer (Spectramax, M2^e^, Molecular devices, US).

### 2.6. Cell Migration Assays

Cell Migration of pretreated cells was tested for chemotaxis and chemokinesis. In this study 1% (v/v) ethanol concentration was used to establish ethanol gradient for chemotaxis and to maintain uniform ethanol concentration for chemokinesis studies.

#### 2.6.1. Transwell Assay (Chemotaxis)

Fibroblasts subjected to respective dosing were harvested. 5×10^4^ cells were then seeded on the upper chamber of the well inserts (24-well plate cell inserts) and incubated for 10 min at 37ºC, 5% CO_2_. The lower chamber was then filled with 600 μl of media with and without 1% alcohol. The upper chamber was kept serum-free, whereas the lower chamber had 10% FBS. After 12 h of incubation, transwell inserts were removed from the plate and the upper chamber was cleaned using a cotton applicator. 600 μl of 70% ethanol was then added to new wells and the inserts were placed to fix the cells for 10 min. To stain the cells for visualization, 600 μl of 0.2% crystal violet was added to fresh wells, and inserts were transferred to it. Staining was done for 10 min at room temperature. The inserts were then washed with distilled water two to three times and visualized under an optical microscope at 10X magnification. 12 images for each group were taken from different areas. The cells were counted in ImageJ software (Schneider et al., 2012).

To understand the role of FBS gradient in cell migration to ethanol, seven different conditions in the Boyden Chamber were checked. These seven conditions are denoted by c1-c7 and have been depicted in a tabular form for better understanding (Table S1 of Supplementary Material 1). For this experiment, ethanol untreated cells (control) were taken and cell density in the upper chamber was kept higher (7 × 10^4^ cells/well).

#### 2.6.2. Random Cell Migration (Chemokinesis)

Cells harvested after ethanol treatment were seeded in a 24-well plate with a cell concentration of 2×10^3^ cells/well. After 4 hrs of seeding one set of treatment groups were subjected to exposure of 1% ethanol (uniform concentration maintained) and the other set, without alcohol. Immediately after ethanol exposure, live imaging of cells was done for 22 h in Zeiss Spinning Disc Confocal Microscope in DIC (Differential Interference Contrast) mode to track real-time cell movement. Migration parameters such as total run length and directionality of non-dividing cells were calculated for 8 h.

### 2.7. Analysis of blebbing

Blebs in cells were characterized from real-time imaging of cells, as described in Section 2.6.2, in terms of the percentage of cells with blebs and cell recovery time. The percentage of cells with blebs was calculated using Eq. 1.

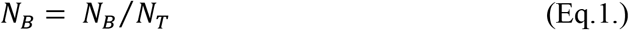

where, *N_B_* is the number of cells with blebs, and *N_T_* is the total number of cells.

Additionally, to determine the blebbing activity or blebbing index, 1 × 10^4^ cells were subjected to live imaging at 40X magnification with 15 min intervals for 2 h. Z-stacking was done to cover all the blebs in all the planes. Blebbing activity was determined using Eq. 2.

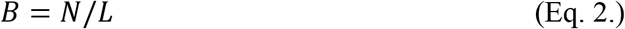

where *B* is the blebbing activity, *N* is the number of blebs on a cell, and *L* is the cell perimeter, at a particular time point.

### 2.8. Analysis of kymograph

Cells harvested after ethanol treatment were seeded in a 35 mm confocal dish at a concentration of 1 × 10^3^ cells/dish and incubated for 12 h. Time-lapse imaging of cells was then done in both normal media and media supplemented with 1% ethanol. Imaging was done in differential interference contrast (DIC) mode at 40X magnification for 30 min with a time interval of 10 s in a confocal microscope (Zeiss Spinning Disk Confocal Microscope, Germany). Kymographs were obtained using lines in ImageJ software and were analyzed for obtaining protrusion and retraction dynamics. At least 3 lines per cell were drawn.

### 2.9. Analysis of cell morphology

5 × 10^3^ cells were seeded in a 12 mm coverslip and incubated for 12 h. The cells were then fixed with 4% paraformaldehyde for 10 min followed by PBS washing. Fixed cells were then permeabilized with 0.1% Triton X (10 min) and stained with Florescene Phalloidin for actin (4 h, 4ºC) and DAPI for nucleus (10 min, RT). Imaging of cells was then done in laser scanning confocal microscopy in 63X oil objective with z stacking (Zeiss LSM 80 Confocal Microscope, Germany). Maximum Intensity Projection (MIP) of the z-stacks was done in Zen Blue software (Zeiss, Germany). Morphological analysis of cells was done in ImageJ software to obtain cell area and length/width (L/W) ratio.

### 2.10. Statistical Analysis

One-way ANOVA (Analysis of Variance) followed by Tukey’s test was used to determine the statistical significance of data obtained from three independent experiments. The test of significance was done in OriginPro 9.1 software (Seifert, 2014). Data from all the experiments have been represented as mean ± standard error of the mean (SE), and p-values less than 0.05, 0.01, and 0.005 were considered statistically significant.

## 3. Results

### 3.1. Cell Proliferation Assay

As seen in Figure 1a, ethanol pre-exposure of 0.01A, 1A, 0.01C, and 1C do not affect cell proliferation. Cell numbers of the 3A and 3C treatment groups are lower than the rest of the groups, with 3C being significantly low. This suggests that ethanol adversely affects the rate of cell proliferation in the 3C pre-treatment group. The anti-proliferative effect of ethanol in fibroblast cells is attributed to inhibition of growth factor-induced signaling pathways for cell growth and disruption of DNA synthesis (Yeo et al., 2000).

**Figure 1:**
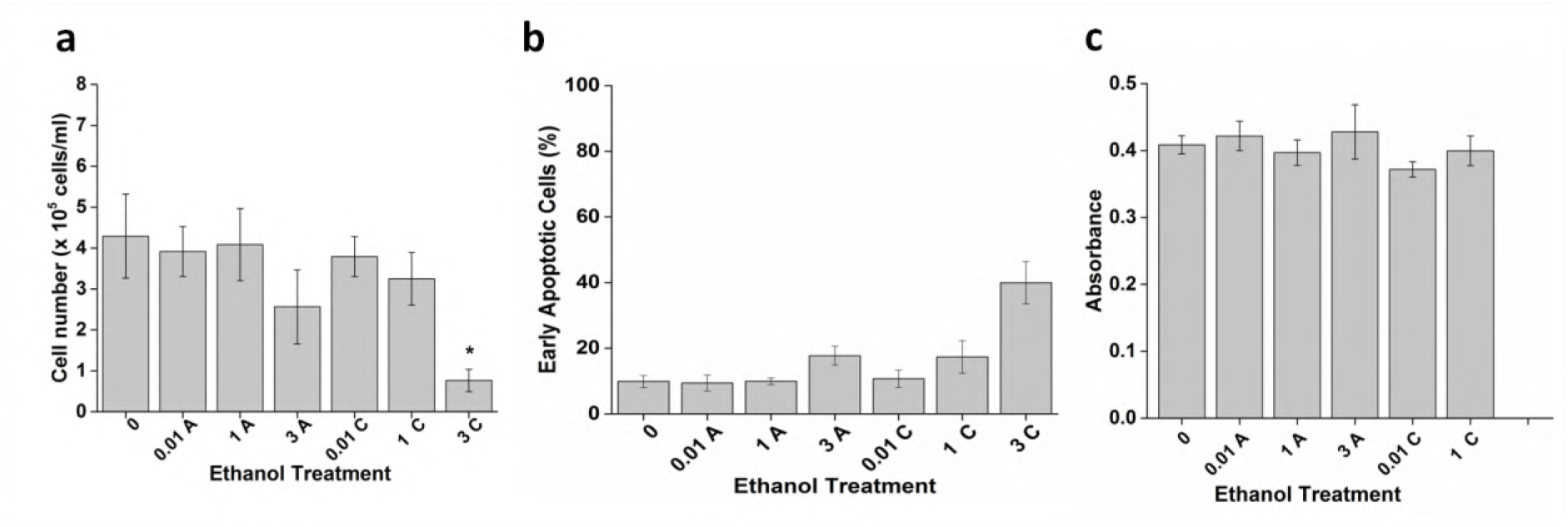
Effect of ethanol treatment on (a) cell proliferation (b) apoptosis, and (c) cell adhesion. The figure depicts that the 3C treatment group has a very low cell count and higher apoptosis after seven days of treatment. Cell adhesion assay suggests that ethanol treatment do not affect the adhesion of cells on the substrate

### 3.2. Cell Apoptosis Assay

Figure 1b indicates that ethanol-induced apoptosis is significant in the 3C pre-treatment group, with around 40% of cells being in the early apoptosis stage. It should also be noted that though not statistically significant, apoptotic cells can also be seen in 3A and 1C pre-treatment groups. This is in line with the fact that cytotoxicity to ethanol depends on both ethanol concentration and exposure time (Tapani et al., 1996). Here, 3A represents short repeated exposure of sub-toxic ethanol concentration of 3%, whereas 1C represents chronic exposure to a moderate ethanol concentration of 1%, thus inducing toxicity. Ethanol toxicity leading to apoptotic death is majorly due to the induction of oxidative stress (Circu and Aw, 2010; Yang et al., 2018).

Owing to the slow proliferation rate and high apoptosis of cells exposed to chronic 3% ethanol exposure, this pre-treatment group (3C) has been excluded from the rest of the studies.

### 3.3. Cell Adhesion Assay

Cell adhesion assay suggests that treatment of cells with ethanol does not affect cell-substrate adhesion. This has been illustrated in Figure 1c, where the measured absorbances are not significantly different for the pre-treatment groups.

### 3.4. Cell Migration Assays

#### 3.4.1. Transwell assay

Figure 2 illustrates the potential of ethanol as a chemotactic agent. In this assay, the migratory behavior of untreated and pretreated cells towards ethanol gradient was evaluated. As clear from the Figure 2b and 2c, both ethanol pre-treated, as well as untreated cells, migrate towards ethanol gradient. However, migration in the case of pretreated cells is higher than that of untreated cells. Fold increase in migration of cells in presence of ethanol gradient with respect to their migration in absence of ethanol gradient has been calculated as 1.3, 2, 2.6, 2.8, 2.5, and 4.1 for 0, 0.01A, 1A, 3A, 0.01C, and 3C pretreatment groups respectively. Moreover, the fold increase in migration of pre-exposed cells as compared to untreated cells (0 or control group) in ethanol gradient is 1.1, 1.5, 1.5, 2.2, and 2.3 for 0.01A, 1A, 3A, 0.01C, and 3C pretreatment groups respectively. Thus migration of cells towards ethanol follows the following trend: untreated < Acute-Periodic < Chronic. This shows that ethanol can not only act as a chemoattractant for cells but also its pre-treatment enhances the chemotactic responsiveness of cells. Dose dependency can be observed in the ability of cells to migrate towards ethanol i.e. chronic exposure of ethanol (0.01C and 1C) led to the higher migrational ability of cells, than acute-periodic exposure (1A and 3A). Furthermore, in absence of ethanol gradient (i.e. no ethanol in the second compartment), migration of cells for untreated and pretreated cells are similar.

**Figure 2:**
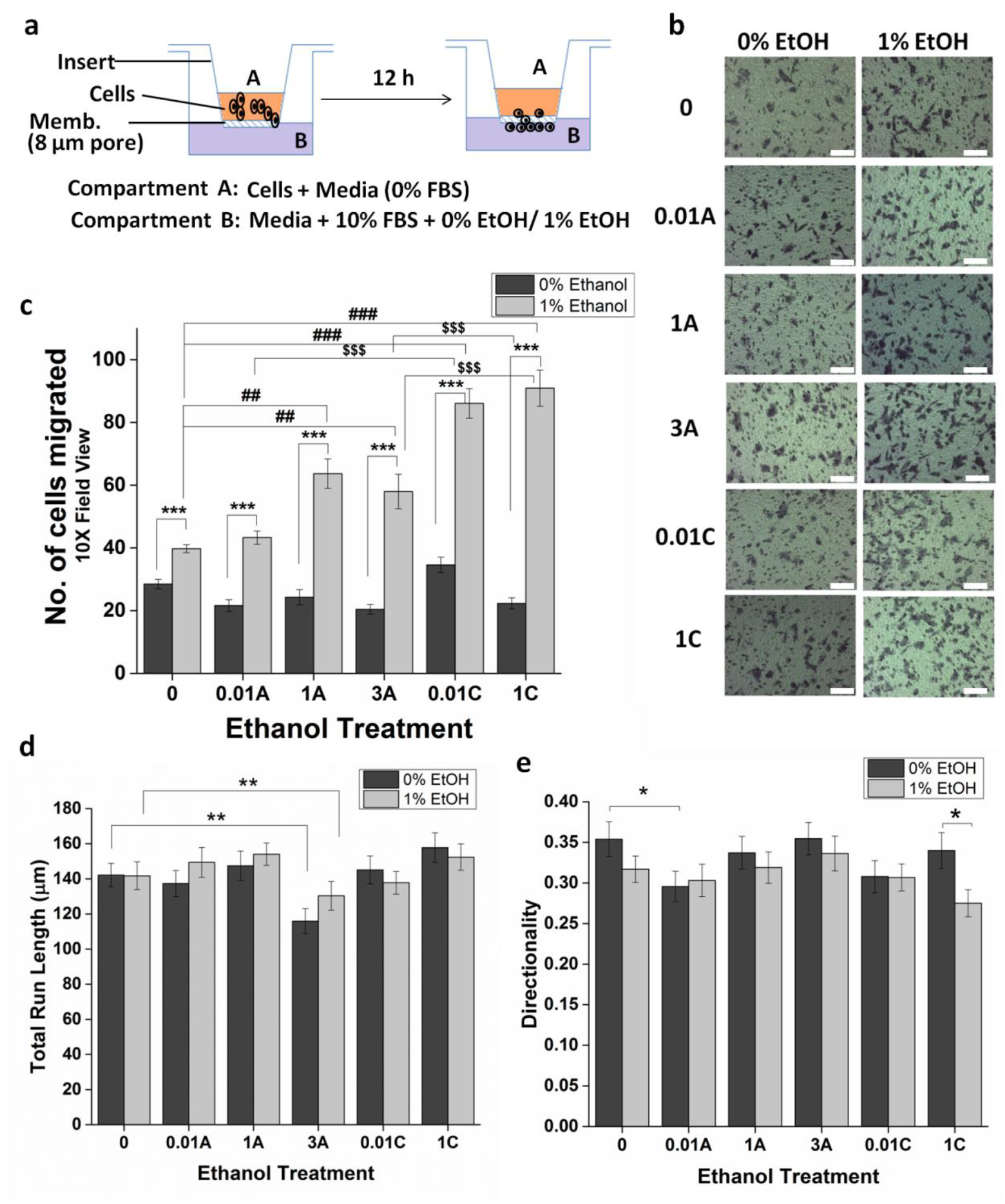
Cell migration assays. (a), (b), and (c) depicts results from the transwell assay. (a) Experimental set up (b) Optical micrographs of the lower side of membrane containing cells (c) Quantitative analysis of migrated cells. Both treated and untreated cells migrate towards compartment B containing ethanol. Migration of pretreated cells towards ethanol is significantly higher than untreated cells. Chronic pre-treatment shows more migration towards ethanol than acute pre-treatment. (d) and (e) depicts results for random cell migration. Effect of ethanol on (d) total run length and (e) directionality. 3A treatment group shows a decrease in total run length. The directionality of the 0.01A treatment group has decreased and 1C shows decreased directionality upon addition of ethanol. Scale bar: 100 μm. ***P < 0.005, **P < 0.01, *P < 0.05, ###P < 0.005, ##P < 0.01, $$$P < 0.005.

It should be noted that ethanol enhances the chemotactic ability of cells only in presence of an FBS gradient. Figure S1 (Supplementary Material 1) represents the result of seven different conditions (c1-c7) to determine the chemotactic ability of fibroblast cells (untreated). As seen in the figure, in absence of an FBS gradient (c2, c4, and c5), there is no migration of cells towards the lower chamber, even if an ethanol gradient of 1% is provided. In presence of both FBS and ethanol gradient, migration of cells is higher when ethanol gradient is maintained with 1% ethanol (c7) rather than 3% ethanol (c6). This might be because 3% ethanol concentration is more toxic. Moreover, migration of cells is higher in presence of both FBS and 1% ethanol gradient (c7), than only FBS gradient (c1).

#### 3.4.2. Random cell migration

Migration of cells in uniform ethanol concentration was tested by time-lapse imaging of untreated and pretreated cells. As soon as 1% ethanol is introduced to the culture plates for maintaining uniform concentration, cells shrink and blebs are formed (discussed in detail in next section), and cell migration stalls for some time, until the cells recover. Figures 2d and 2e illustrate two different parameters calculated i.e. total run length and directionality. The results suggest that except for 3A, pre-treatment does not affect cell velocity in the presence or absence of ethanol i.e. total run length covered by cells in 8 h is the same as that of untreated cells. 3A pre-treatment group covers a shorter total run length. As far as directionality is concerned, pre-treatment group 0.01A shows low directionality, i.e. tendency of random walk is more in this group in both presence and absence of ethanol. Overall, the migration ability of pre-treated cells is not significantly affected during chemokinesis. These results suggest that migration of cells towards ethanol is externally regulated and pre-exposure to ethanol does not cause a significant change in intrinsic cell directionality, which is otherwise known to affect chemotaxis (Petrie et al., 2009).

### 3.5. Analysis of blebbing

Figure 3a and 3b depicts that revival of cells from ethanol insult occurs in three distinct stages: (a) shrinking, (b) blebbing, and (c) recovery. As soon as ethanol is added to the cells (both pre-treated and untreated), cells shrink, and small membrane blebs appear on the cell surface (Figure 3a2). Thereafter, the small blebs are replaced by large dynamic blebs (Figure 3a3), due to actomyosin contraction and an increase in hydrostatic pressure (Charras et al., 2008). Interestingly, these two stages of cell morphology are also seen in cells undergoing apoptosis (Charras, 2008). However, in this case, the blebs eventually retreat, cells spread and return to normal morphology (Figure 3a4). This suggests that cells evade lysis and adapt to the new environment by the means of cellular blebbing. The role of blebbing in protection against cell injury has also been identified earlier (Babiychuk et al., 2011).

**Figure 3:**
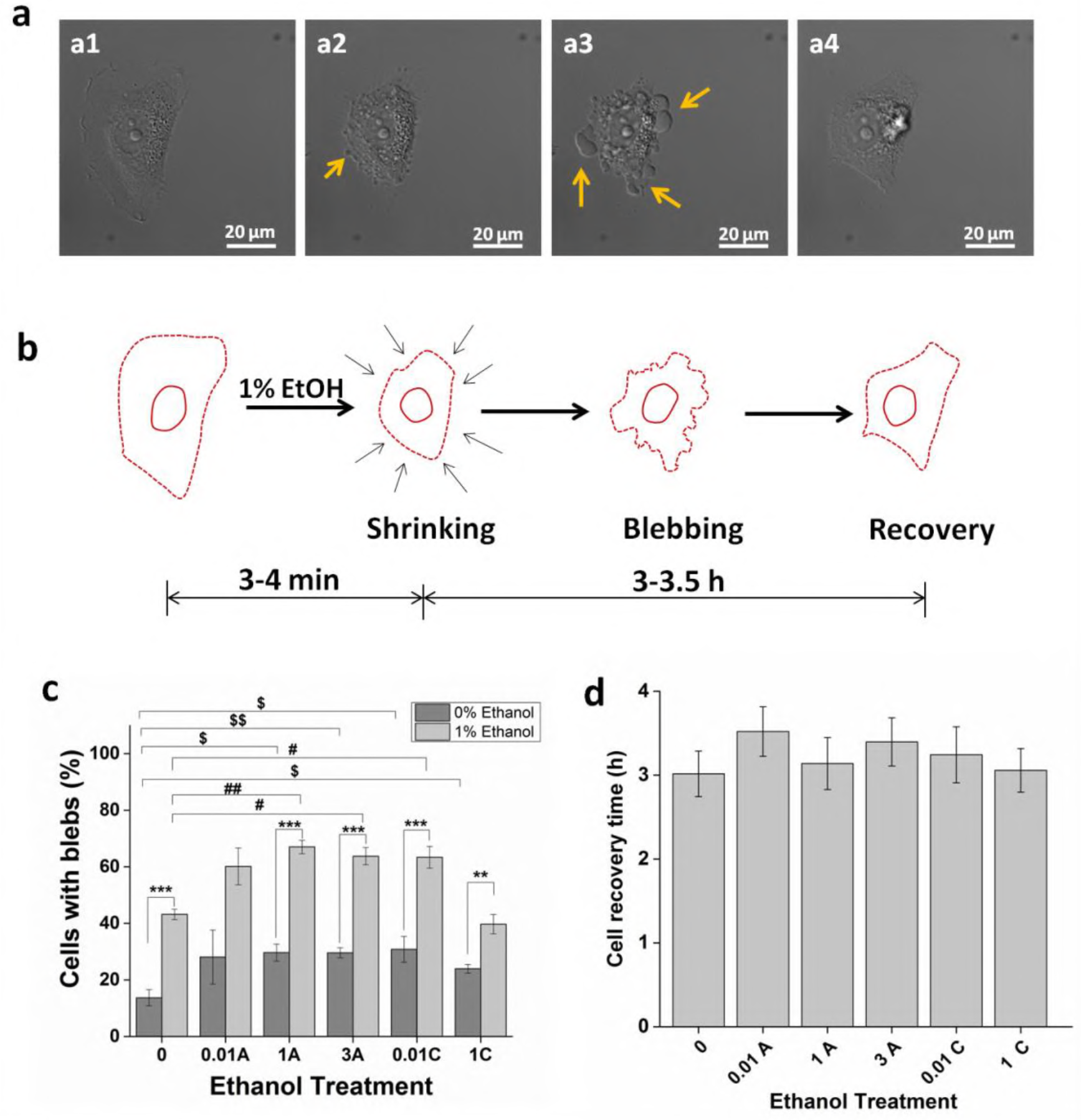
Analysis of blebs. (a) DIC images, and (b) schematic diagram for stages of cell morphology upon ethanol exposure. The addition of 1% ethanol first leads to shrinking (a2), then dynamic blebbing (a3), and finally cell recovery (a4). Yellow arrows depict blebs. (c) Percentage of cells with blebs. Exposure to 1% ethanol increases the percentage cell blebbing in pretreated and untreated cells. However, the percentage cell blebbing in pretreated cells is more than that of untreated cells except for 1C. (d) Cell recovery time for all the pre-treatment groups is the same. ***P < 0.005, ##P < 0.01, #P < 0.05, $$P < 0.01, $P < 0.05.

As observed in Figure 3c, pre-treatment of fibroblasts led to the appearance of blebs on the cell surface. Around 2 fold increase in the percentage of cells with blebs is observed in ethanol pre-treated cells. Moreover, upon exposure to 1% ethanol (for maintenance of uniform ethanol concentration), the percentage of cells with blebs increases for both pretreated and untreated cells.

However, the percentage of cells with blebs in pre-treatment groups is significantly higher than untreated cells, except for the 1C treatment group. Furthermore, a fold increase in the percentage of cells with blebs in presence of 1% ethanol with respect to the absence of ethanol is calculated as 3.1, 2.1, 2.3, 2.2, 2.1, and 1.7 for 0, 0.01A, 1A, 3A, 0.01C, and 1C respectively. Thus, cells previously unexposed to ethanol, show the highest change in the number of cells with blebs in presence of 1% ethanol, whereas pre-exposing cells with 1% ethanol chronically, show the lowest change upon further addition of 1% ethanol. This indicates that untreated cells show the highest resistance to change in the environment. However, regardless of treatment groups, the blebs are short-lived and retreat from the cell surface eventually in ̴ 3 h, as observed in Figure 3d.

Figure 4 depicts the blebbing activity of cells for 2 h. It indicates the life of blebs upon ethanol exposure. The figure indicates that the blebbing activity upon ethanol exposure follows a bell-shaped curve with the highest activity after 15 min of exposure. However, the activity at 15 min is low for the acute-periodic pre-treatment group and the highest activity is for the untreated and 0.01C treated group.

**Figure 4:**
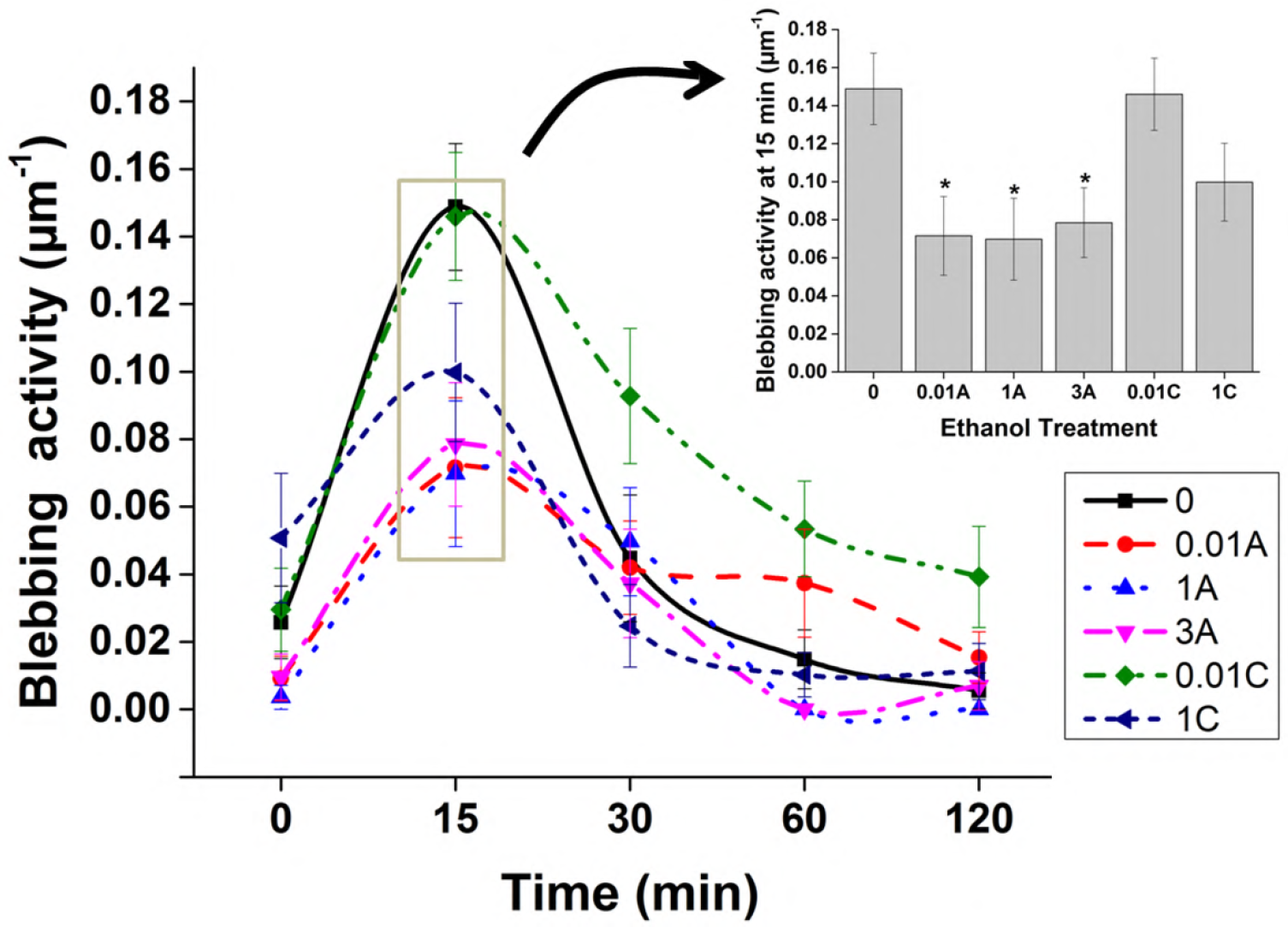
Analysis of blebbing activity. (a) Blebbing activity at different time points (b) Blebbing activity at 15 min time point. The blebbing activity upon ethanol exposure over a period of time follows a bell-shaped curve, with maximum blebbing activity at 15 min. *P<0.05.

### 3.6. Analysis of protrusion dynamics

Analysis of kymographs from time-lapse imaging has been depicted in Figure 5. Parameters such as protrusion velocity, retraction velocity, and protrusion frequency have been calculated from kymographs. Figures 5b and 5c depict protrusion and retraction velocities of pre-treated and untreated cells. These figures indicate that both protrusions, as well as retraction velocities of ethanol pretreated cells, are more than that of untreated cells. Also, exposure to 1% ethanol further increases these velocities of both pretreated and untreated cells.

**Figure 5:**
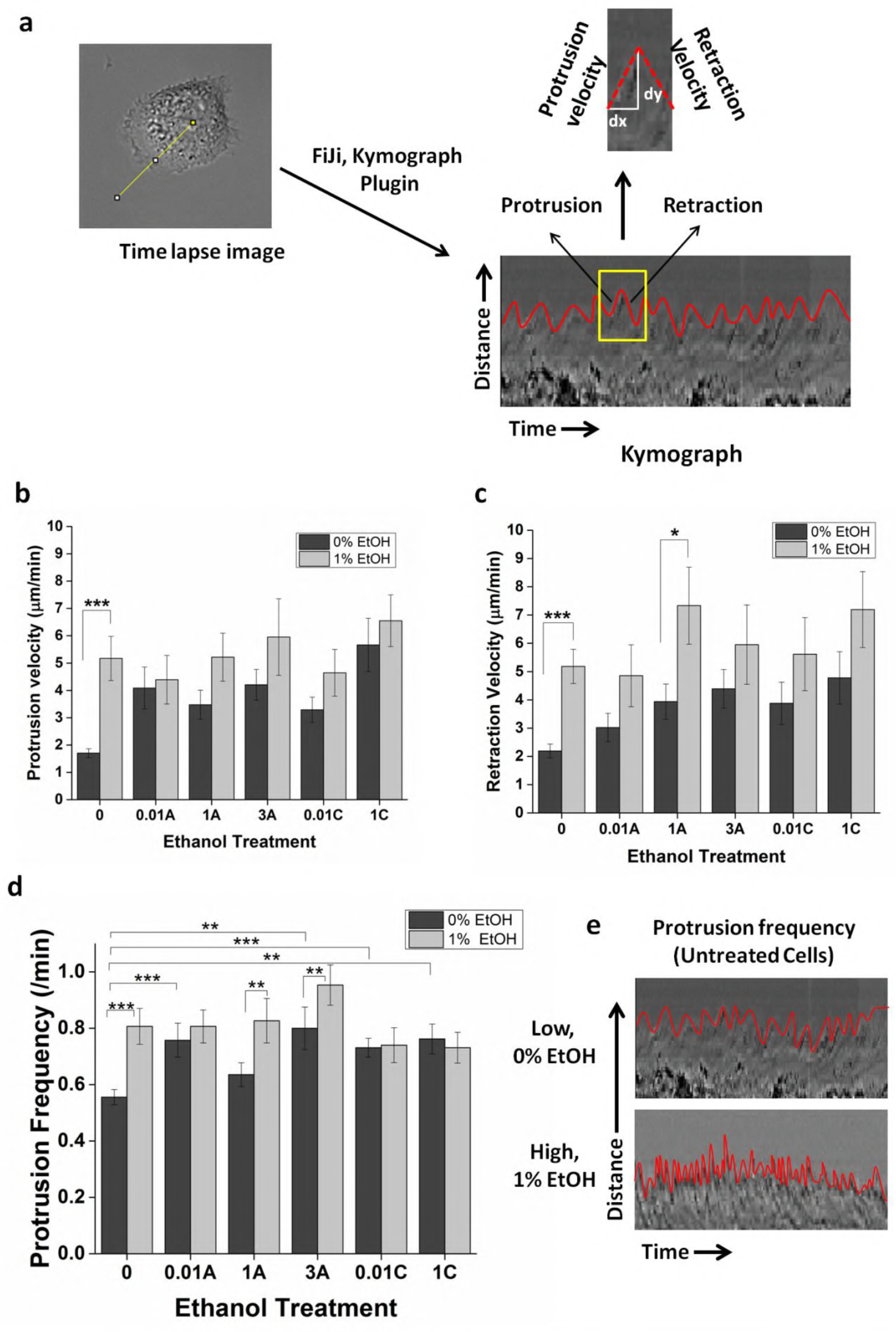
Analysis of protrusion dynamics. (a) Scheme for obtaining kymograph. A line drawn from the center of the cell (time-lapse image) towards the cell periphery is used to obtain the kymograph. The x-axis of the kymograph represents time, whereas the y-axis represents distance. Corresponding protrusion and retraction velocities are then calculated. (b) Protrusion Velocity, and (c) Retraction Velocity. Protrusion and retraction velocity of pretreated cells is more than untreated cells in absence of ethanol. In presence of ethanol both the velocities increases for pretreated as well as untreated cells. (d) Protrusion Frequency. Protrusion frequencies of pretreated cells are more than that of untreated cells. However, exposure to 1% ethanol increases the protrusion frequencies of untreated as well as acute-periodic treated cells. (e) Kymographs representing low and high protrusion frequencies. Here, untreated cells exposed with and without ethanol have been presented as an example.

Figures 5d and 5e illustrate the protrusion frequencies of pre-treated and untreated cells. Protrusion frequency refers to the number of protrusions per minute. Similar to protrusion and retraction velocities, protrusion frequencies of pretreated cells are also more than that of untreated cells. However, upon 1% ethanol exposure, protrusion frequencies of only untreated cells and acute-periodic treated cells increases. These results suggest that cell membrane activity increases upon pre-treatment and the addition of ethanol further makes it highly dynamic, possibly due to the development of blebs.

### 3.7. Cell morphology analysis

Figure S2 (Supplementary Material 2) depicts fluorescence micrographs of ethanol-treated cells. Quantitative analysis of cell morphology has been done in terms of L/W ratio (length/width) and cell surface area. As seen in Figures S3a and S3b, ethanol treatment of 0.01A has resulted in a lower L/W ratio and higher cell surface area. The hypertrophy of 0.01A cells explains the lower directionality of the cells in the chemokinesis setup. Other pre-treatment groups do not significantly affect these parameters. Moreover, acute-periodic pre-treated cells undergo the least reduction in cell size upon subsequent 1% ethanol exposure Figure S3c.

## 4. Discussion

This study illustrates the effect of ethanol and its pre-exposure on eukaryotic cell migration, specifically fibroblasts. The idea is to determine if alcohol-seeking behavior is exhibited even at a cellular level. Thus, the potential of ethanol as a chemotactic agent has been evaluated, with a focus on how pre-exposure affects it. In this study, two different dosing strategies were used to pre-treat fibroblast cells (acute-periodic and chronic) and their migrational dynamics were evaluated. Both chemotaxis and chemokinesis were tested along with the protrusion dynamics of cells.

This study brings out three significant observations pertaining to ethanol-induced directed cell migration: (a) fibroblast cells (both pre-exposed and unexposed) show affinity towards ethanol, (b) ethanol affinity of pre-exposed cells is significantly greater than unexposed cells, and (c) ethanol affinity of cells with chronic pre-exposure is greater than cells with acute-periodic exposure. These observations suggest that ethanol not only acts as a potent chemoattractant for fibroblast cells but also its pre-exposure enhances the cells’ chemotactic responsiveness. Also, the strength of pre-exposure determines the migratory behavior towards ethanol. This is in agreement with the general fact that alcohol consumption patterns and the development of alcoholism are interrelated (Wackernah et al., 2014). Interestingly, among other drugs of addiction, pre-exposure to nicotine and morphine is known to enhance chemotactic responsiveness of human neutrophils (Totti et al., 1984) and microglia (Horvath and DeLeo, 2009) to specific chemoattractants respectively. Moreover, the migratory response elicited by these drugs is concentration-dependent and increases with an increase in strength of pre-exposure, as also observed in this study. However, chemotaxis to recreational drugs per se is an unexplored area of research.

The ability of cells to sense ethanol gradient may be indicative of the presence of G-protein coupled receptors on the cell surface, specific for ethanol, which in turn elicits one or the other known signaling cascades responsible for directed cell migration (Devreotes and Janetopoulos, 2003; Petrie et al., 2009). As far as enhanced chemotactic responsiveness is concerned, we hypothesize that pre-exposure to alcohol up-modulates the expression of surface receptors, thus affecting the chemotactic activity. The hypothesis is drawn from the fact that chemotaxis can be regulated by the alteration of chemoattractant receptors (Franitza et al., 2002; Seely et al., 2002). Increased chemotaxis to one chemoattractant upon pre-exposure to another has been reported in eosinophil cells (Schratl et al., 2006). Moreover, an altered protrusion dynamics of pre-exposed cells might also contribute to the development of seeking behavior in cells (Diz-Muñoz et al., 2016).

When 1% ethanol is introduced to the cellular environment of pre-treated cells to maintain ethanol homogeneity, cells initially shrink. The shrinkage of cells is caused by plasma membrane fluidization (Goldstein, 1986) and cytoskeletal disaggregation (Iwata et al., 2011). This is followed by cell blebbing, which is a highly dynamic process. The protrusion/retraction velocities and protrusion frequencies of the cell membrane increases. A bell-shaped curve is obtained for the blebbing activity of the cells, with a maximum number of blebs seen at around 15 min (T_max_) of ethanol exposure, that almost disappear after 2 h of exposure. Complete cell recovery and regain of morphology occur after 3 h. Cell migration is then initiated. Blebs are generally considered as a hallmark of apoptosis (Barros et al., 2003), however, it also fulfills a variety of other functions in cells viz. cell division (Charras et al., 2008), migration (Diz-Muñoz et al., 2016), and recently discovered the function of damage control (Babiychuk et al., 2011). Since the cells in consideration survive, the blebs formed are non-apoptotic. Only non-dividing cells have been considered for the study and migration is stalled during the lifetime of the blebs, thus the possibility of bleb-driven damage control of cells may come into play here. Moreover, ethanol injury causes an increase in cytoplasmic Ca^2+^ concentration (Del Castillo‐Vaquero et al., 2010; Gandhi and Ross, 1989) and an increased Ca^2+^ concentration initiates bleb-driven damage control machinery of cells (Babiychuk et al., 2011). It should also be noted that mild pre-exposure to ethanol resists shrinkage (0.01A, 1A), with lower blebbing activity at T_max_ (0.01, 1A, and 3A). This may be an incidence of pre-conditioning hormesis, where prior exposure to mild doses of toxic compound helps the system to better adapt upon subsequent high dose exposure (Calabrese, 2016).

## 5. Conclusions

Overall, this study establishes ethanol as a chemoattractant for fibroblast cells, highlighting the fact that seeking behavior can be traced down to even the cellular level of organization. Moreover, bleb-driven machinery of cell protection to survive and adapt in presence of ethanol is also a significant finding of this study. This cellular approach adds a new dimension to addiction research, which can be further explored to understand the underlying genetic variability that contributes to ethanol addictive behavior. The study thus widens our understanding of ethanol-seeking behavior which was earlier confined to only the higher organisms and usually attributed to neural networks of the central nervous system.

## 7. Acknowledgments

The authors are thankful to Central Facility, Industrial Research and Consultancy Centre (IRCC), IIT Bombay, for the use of Spinning Disk Confocal Microscope.

## 8. Author Contributions

Neelakshi Kar performed all the experiments and wrote the main text of the manuscript. Jayesh Bellare is the project advisor and helped with experiment design, data analysis, and drafting of the manuscript. All authors approved the final version of the manuscript.

## 10. Data Availability

All data generated or analyzed during this study are included in the article.

