## Supplementary Material 1 for "Ethanol pre-exposure enhances alcohol-seeking behavior at cellular level by chemoattraction and exhibits bleb-driven cellular stress response in uniform ethanol concentration"

**Effect of FBS and ethanol gradient on cell migration**

Table S1: Different experiment conditions for ethanol and FBS gradient

| **Condition** | **Compartment A (Upper chamber)** | **Compartment B**  **(Lower chamber)** |
| --- | --- | --- |
| c1 | Media + Cells | Media + FBS |
| c2 | Media + Cells + FBS | Media + FBS |
| c3 | Media + Cells + FBS | Media |
| c4 | Media + Cells + FBS | Media + FBS + 1% EtOH |
| c5 | Media + Cells | Media + 1% EtOH |
| c6 | Media + Cells | Media + FBS + 3% EtOH |
| c7 | Media + Cells | Media + FBS + 1% EtOH |

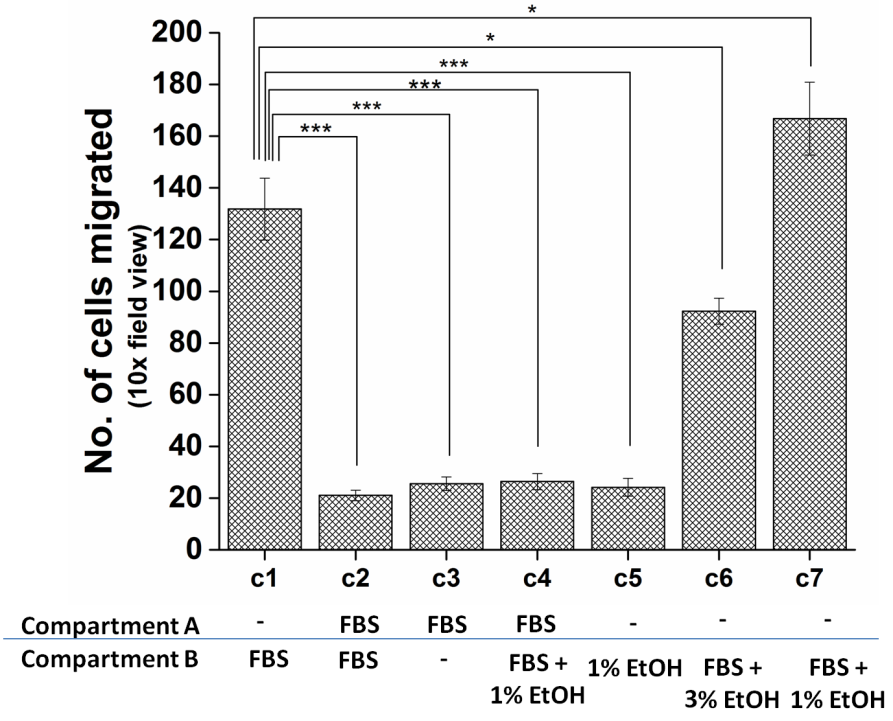

Figure S1: Quantitative analysis of transwell assay showing importance of FBS gradient. The figure shows 7 different conditions of ethanol and FBS gradients. The figure shows that ethanol enhances the chemotactic ability of cell only in presence of FBS gradient (c7).
