## Supplementary Material 2 for "Ethanol pre-exposure enhances alcohol-seeking behavior at cellular level by chemoattraction and exhibits bleb-driven cellular stress response in uniform ethanol concentration"

**Effect of ethanol pre-treatment on cell morphology**

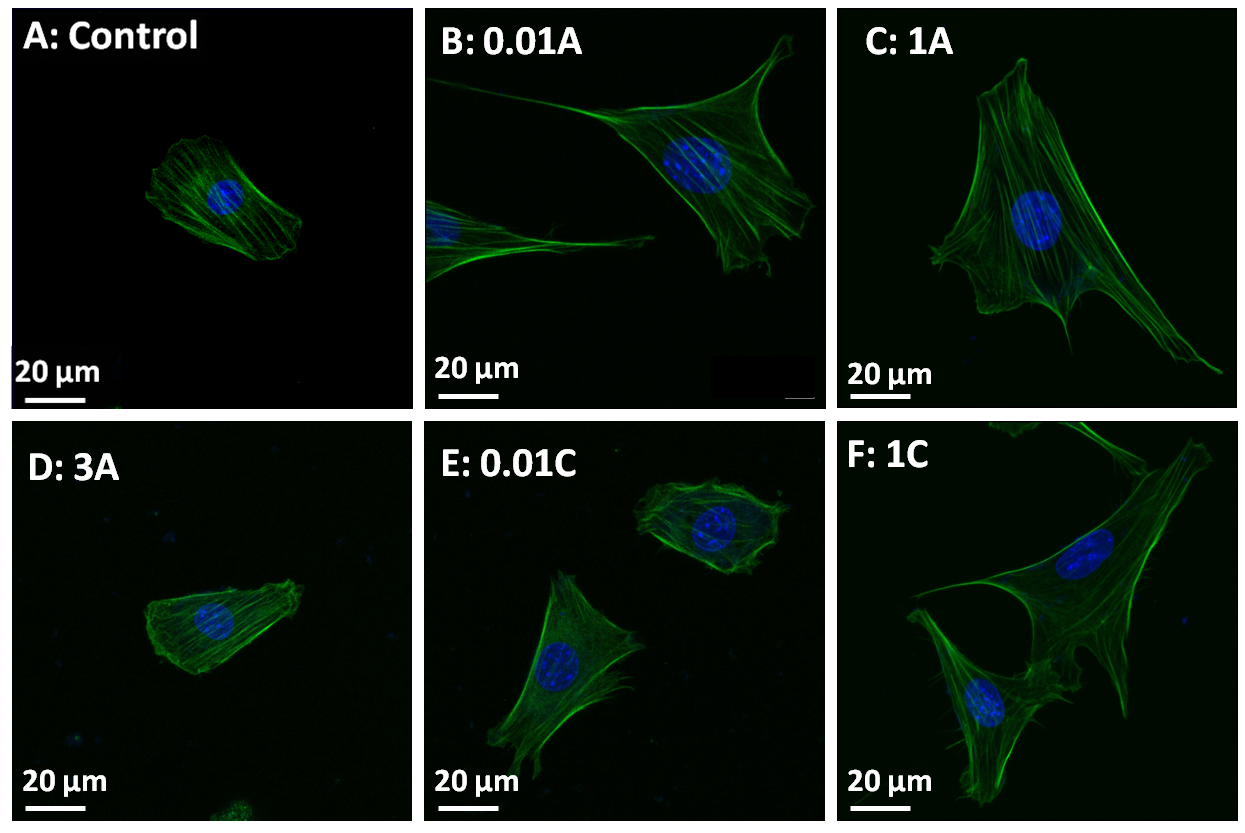

Figure S2: Fluorescence micrographs of ethanol treated cells. Cells stained with FITC-Phalloidin and DAPI. (A) Control (B) 0.01A (C) 1A (D) 3A (E) 0.01C (F) 1C

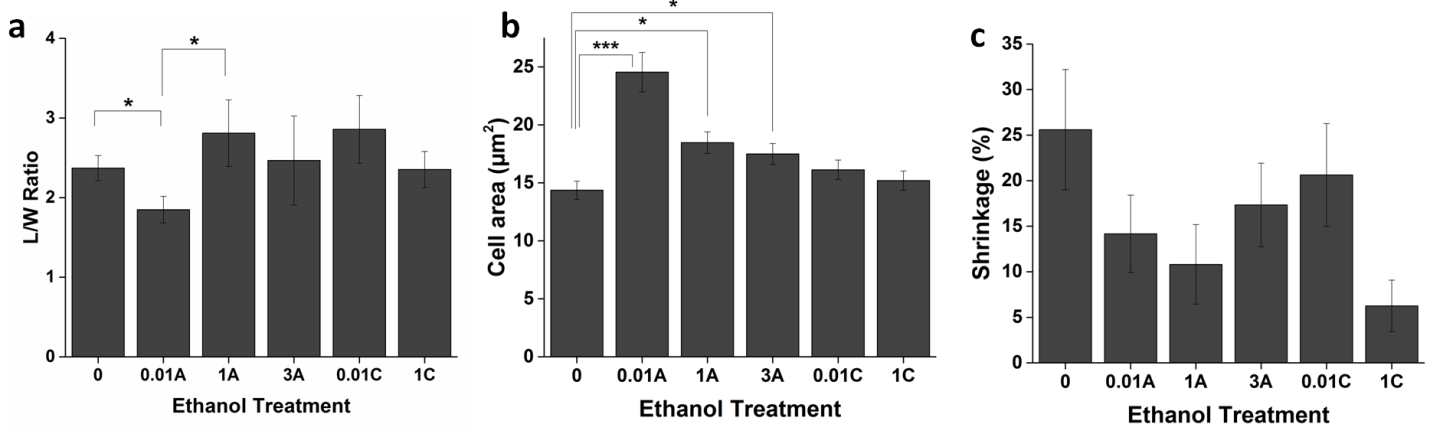

Figure S3: Quantitative analysis of cell morphology. (a) Length/Width ratio (b) Cell surface area (c) Percentage shrinkage of cells upon ethanol exposure. ***P<0.005, **P<0.01 *P<0.05
